# Trophic plasticity of the highly invasive topmouth gudgeon (*Pseudorasbora parva*) inferred from stable isotope analysis

**DOI:** 10.1101/617688

**Authors:** Matteo Rolla, Sofia Consuegra, Carlos Garcia de Leaniz

## Abstract

A wide trophic niche and high trophic plasticity are often invoked to explain the successful establishment of many aquatic invaders, but there is little information regarding the diet of most invasive fish in European waters. We combined stomach content and stable isotope analysis (SIA) of ^13^C and ^15^N to examine the trophic niche of the highly invasive topmouth gudgeon (*Pseudorasbora parva*) in four contrasting ponds and reservoirs in South Wales. Marked differences in diet and trophic position were found among neighbouring systems only a few kilometres apart. The most diverse diet was found in ponds with the fewest number of inter-specific competitors, and resulted in topmouth gudgeon having the highest condition factor, the widest variation in δ^13^C and δ^15^N values, and the highest carbon content, typical of generalist feeders. In contrast, topmouth gudgeon that coexisted with other cyprinids, were much more likely to have empty stomachs and relied almost exclusively on plant seeds, resulting in fish having a poor condition factor and low trophic position. Comparisons with other trophic studies indicate that variation in isotope values among neighbouring sites can exceed variation across continents, making it difficult to predict the diet or trophic impacts of the invasive topmouth gudgeon. Given the importance of obtaining reliable data on trophic position for risk assessment, our study shows that the use of SIA could be used to prioritize control and eradication measures that take into account trophic plasticity.

## Introduction

The topmouth gudgeon (TMG, *Pseudorasbora parva*) is a highly invasive fish native of Asia that has become one of the worst aquatic invaders in Europe due its potential impact on native biodiversity (Britton et al. 2007; Pinder et al. 2005). TMG possesses many ecological traits that make it a successful invader, including short generation time, high fecundity, and substantial phenotypic plasticity (Gozlan et al. 2010). The species can also grow rapidly (Adrović, Skenderović 2007; Kapusta et al. 2008; Ye et al. 2006) and mature fast (Pinder et al. 2005), especially when living at high density (Britton et al. 2008; Britton et al. 2007).

During the first few years of the invasion, TMG appear to be composed mainly of young-of-the-year, up to 50 mm in length, although the species can reach up to 120 mm (Adrović, Skenderović 2007; Kapusta et al. 2008; Ye et al. 2006). The species displays substantial variation in the timing of spawning, depending on water temperature (Yan, Chen 2009). Its diet can be wide and, although previous studies had described it as mostly omnivorous (Weber 1984; Xie et al. 2001), it can also adopt a strict planktivorous diet in some cases (Asaeda et al. 2007; Priyadarshana et al. 2001). Depending on prey availability, TMG can feed on chironomid larvae (Wolfram-Wais et al. 1999), but also on the eggs and larvae of other fish, which may pose a threat to native species (Stein, Herl 1986; Xie et al. 2000). The preferred prey size appears to be size dependent, as observed in a comparative pond study where the smaller size class (20–25 mm) fed exclusively on cladoceran zooplankton, the bigger size classes (35–60 mm) fed mainly on chironomid larvae, and the intermediate size classes fed on a mixture of the two preys (Declerck et al. 2002). The diet of topmouth gudgeon can also change seasonally, shifting from chironomid larvae in spring and summer to ostracods in winter (Xie et al. 2000). Where introduced, topmouth gudgeon has been found to compete for food with native species in Great Britain (Britton et al. 2010), Belgium (Declerck et al. 2002), and Poland (Witkowski 2002) which can lead to depressed growth rates, reduced reproduction outputs and shifts in the trophic position of native fish (Britton et al. 2010). When other invasive species are present, the topmouth gudgeon’s plasticity in food selection allows it to shift its trophic niche and co-exist with other invaders (Jackson, Britton 2014).

The ecological impacts caused by the TMG are often visible after only a few years from their introduction. For example, Britton et al. (2007) found that three years after its introduction into a lake, TMG was the dominant fish species within the <70 mm size class, and within four years it was the only species successfully recruiting in the lake. The presence of *P. parva* can cause substantial damage to freshwater fisheries, as they can rapidly dominate fish communities and alter trophic web structure. A comparison of ponds with and without TMG showed that the presence of this invasive species resulted in depressed growth rates (and also production) of resident fish, and also a shift in trophic levels (Britton et al. 2010). Another relevant threat posed by TMG is its ability to carry non-native pathogens such as the intracellular parasite *Sphaerothecum destruens* that can become lethal for salmonids (Andreou et al. 2011; Gozlan et al. 2005). For these reasons, it is feared that the spread of TMG could result not just in a significant loss of native river biodiversity, but also on substantial economic losses caused by damage to sport fisheries and salmon farming alike (Gozlan et al. 2005).

TMG was first recorded in Great Britain in 1996, in a pond in Southern England (Domaniewski, Wheeler 1996). It is thought that the first introduction may have occurred a decade earlier, in the mid 1980’s, when TMG originating from Germany were delivered to an ornamental aquaculture facility in Hampshire, and subsequently became established in Lake Tadburn (Gozlan et al. 2002). Within few years, TMG had spread rapidly and has now been reported in 32 locations across England and Wales, most of which are lentic systems, and 10 of which are connected to major catchments (Britton et al. 2007; Britton et al. 2010). Due to its remarkable speed of colonization, and the potential for major ecological and economic impacts, TMG has been classified as highly impactive by the UK Technical Advisory Group on the Water Framework Directive (Panov et al. 2009) and is the subject of costly eradication programmes (Robinson et al. 2019).

Stable isotope analysis (SIA) can be used to study food webs (Vander Zanden et al. 1999), based on the fact that the ratios of heavy to light isotopes in animal tissues are often affected by diet (DeNiro, Epstein 1978, 1981). Carbon and nitrogen isotopes (δ^13^C and δ^15^N, respectively) are the ones most commonly used in freshwater biology, as they are related to the sources of primary productivity (δ^13^C) and to trophic level (δ^15^N). As trophic markers, SIA can be used at different ecological levels, to study variation among individuals, species, and also among communities (Newsome et al. 2009; Whitledge, Rabeni 1997). They can be used to examine trophic position (Cherel et al. 2008; Roth et al. 2006), animal migrations (Cherel et al. 2007; McClellan et al. 2010), impacts of invasive species (Nilsson et al. 2012; Vander Zanden et al. 1999) the contribution of allochthonous versus autochthonous food resources in aquatic ecosystems (Solomon et al. 2011; Venarsky et al. 2014).

Here we employed SIA of nitrogen and carbon, complemented with analysis of stomach contents, to investigate variation in the trophic ecology of topmouth gudgeon in four contrasting freshwater habitats in South Wales. Our aims were two: (1) to describe the trophic niche of this highly invasive species to gain a better understanding of the potential for interference competition with native fish, and (2) to assess the value of SIA for monitoring diet plasticity of a rapidly expanding invasive species in different colonized habitats.

## Methods and Materials

### Origin of samples

We analysed 117 topmouth gudgeon originating from four contrasting water bodies in South Wales (Figure 1, Table 1): two small decorative ponds with few or no species other than eel (Turbine Pond, TUR, n= 27; Dyfatty pond, DYF, n = 28), an eutrophic pond used for recreational fishing regularly stocked with eight species of coarse fish (Sylen Lake, SYL, n = 30), and a larger, cooler reservoir used for water supply and stocked with salmonids (Upper Lliedi Reservoir, LLI, n = 32). Sylen Lake, Turbine pond, and Dyfatty pond were fished between November 2012 and February 2013 while the Upper Lliedi Reservoir (LLI) was fished in June 2018. Specimens were kept frozen until analysis.

**Figure 1.**
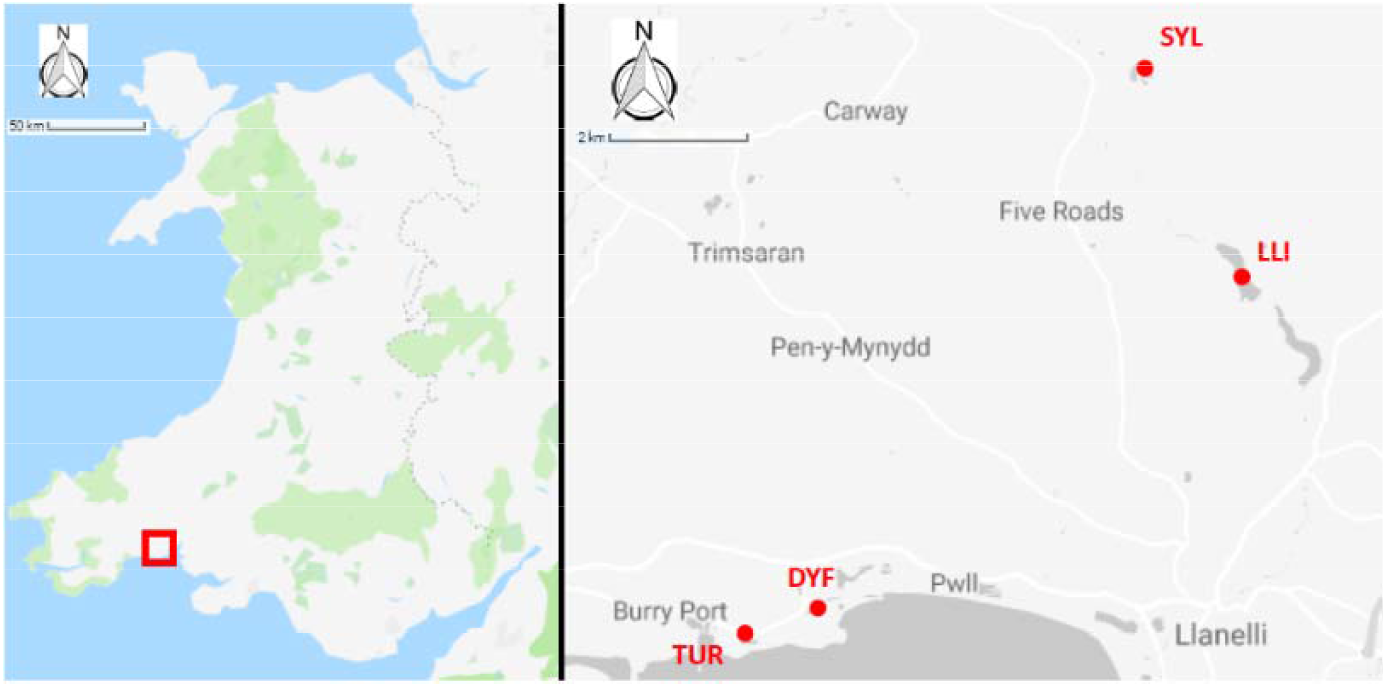
Location of study sites.

**Table 1.** Characteristics of the four study sites sampled for trophic position of topmouth gudgeon (*Pseudorasbora parva*) in South Wales.

| Site | Pond Name | Site GPS | Surface | Altitude (MASL) | Sampling Period | Use | Other fish species | First topmouth gudgeon record | Number of fish sampled |
| --- | --- | --- | --- | --- | --- | --- | --- | --- | --- |
| LLI | Upper Lliedi | SN 5109 0459 | 0.13 km<sup>2</sup> | 86 | June 2018 | Public water supply<br>Recreational angling | Brown trout<br>Rainbow trout<br>Others | NA | 32 |
| SYL | Sylen | SN 49914 07018 | 16,200m<sup>2</sup> | 144 | November 2012 | Fishery<br>Recreational angling | Carp<br>Tench<br>Bream<br>Roach<br>Chub<br>Rudd<br>Bream<br>Goldfish | 2005 | 30 |
| TUR | Turbine | SN 45021 00334 | 10,600m<sup>2</sup> | 7 | February 2013 | Decorative | Eel | 2001 | 27 |
| DYF | Dyfatty | SN 45829 00627 | 3,800m<sup>2</sup> | 4 | February 2013 | Decorative | Eel<br>Perch<br>Rudd<br>Roach | 2001 | 29 |

### Morphometric measurements

We took paired measurements of total length (from the tip of the mouth to the tip of the tail) and weight of TMG from one site (LLI) before (fresh) and after they had been frozen to derive a regression equation to estimate fresh body size from frozen specimens collected from the other three populations. The regression equations were:

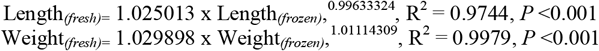

We calculated Fulton’s condition factor (K) from total length (TL) and weight (WT) measurements (K = (WT * 100) / TL^3^) to derive an index of body condition (Bolger, Connolly 1989) used mixture analysis in PAST 3.2.2 (Hammer et al. 2001) to estimate the age composition from length frequency data.

### Stomach content analysis

For stomach content analysis, frozen samples were slowly thawed and then transferred to 30%, 50% and 70% ethanol solutions. We analysed stomach contents of fish in three ponds with a stereo microscope (Nikon SMZ1270) at 12.7x (0.63 - 8x) magnification. The vast majority of prey consisted of zooplankton (67%) that we classified at family level. Insect parts (1%), seeds (24.1%) and plant material were also present, but these could not be identified. We counted every food item and measured the size (longest axis) of a subsample of food items to obtain an average size for each prey category.

### Stable isotope analyses

We obtained 1-2 mg of white muscle from the caudal part of the fish, above the lateral line, and froze dried the samples for 24 hours before analysis. Analyses of %C, %N, δ^13^C, and δ^15^N were conducted by Isotope Ratio Mass Spectrometry (EA-IRMS) at Iso-Analytical Ltd (Marshfield Bank, Crewe CW2 8UY, UK) using a Europa Scientific 20-20 IRMS Elemental Analyser and combusted at 1000 °C in the presence of oxygen using IA-R068 as reference material (soy protein, δ^13^C_V-PDB_ = −25.22 ‰, δ^15^N_AIR_ = 0.99 ‰). The following standards were run as quality controls: IA-R068, IA-R038 (L-alanine, δ^13^C_V-PDB_ = −24.99 ‰, δ^15^N_AIR_ = −0.65 ‰), IA-R069 (tuna protein, δ^13^C_V-PDB_ = −18.88 ‰, δ^15^N_AIR_ = 11.60 ‰) and a mixture of IAEA-C7 (oxalic acid, δ^13^C_V-PDB_ = −14.48 ‰) and IA-R046 (ammonium sulfate, δ^15^N_AIR_ = 22.04 ‰) were run as quality control samples during analysis and were calibrated against inter-laboratory standards distributed by the International Atomic Energy Agency, Vienna. Repeatability of isotope values in samples analysed in duplicate (20%) was high (Cohen’s weighted kappa coefficient: δ^13^C = 0.99; δ^15^N = 0.98; C:N = 0.94) indicating that SIA results were precise. Duplicates were then averaged before numerical analysis. The carbon to nitrogen ratio (C:N) ranged between 3.13 and 3.81, but was significantly below 3.5 (U95CI = 3.35), so lipid correction normalization was not deemed necessary (Post et al. 2007).

### Statistical analysis

We used R 3.3 (R Core Team, 2017) for all analysis except mixture age analysis that was carried out using PAST (Hammer et al. 2001). We transformed % carbon and % nitrogen content with the arcsine transformation to make the data suitable for linear model analysis. We analysed nitrogen and carbon content %, δ^13^C, δ^15^N and C:N ratio as function of fish size, condition factor and pond (location), and used the *dredge* and *anova* functions in the MuMIn R package for model selection. We calculated the Shannon diversity index (*H*) to estimate the taxonomical diversity of the food items present in fish stomachs. Quadratic weights were applied to give a greatest emphasis to large differences between scores. GLM binary logistic regression was used to model the occurrence of empty stomachs (yes or no) as a function of fish size, condition factor and pond type. We performed an analysis of similarities (ANOSIM) with the *vegan* R package (Oksanen et al. 2013) to test for differences in diet composition among locations, merging together the less frequent food classes (unidentified, insect parts and *Cyclops*). Differences in the Shannon index of diversity (*H*) were assessed by bootstrapping (1,000 permutations) to obtain means and 95 confidence intervals (Gardener 2014), we then used the Hutcheson t-test to compare the diversity among sites (Hutcheson 1970).

Trophic breadth for each pond was calculated using Levin’s measure of niche breadth:

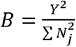

where

B = Leviny’s measure of niche breadth

N_j_ = Number of individuals using prey_j_

Y = Σ N_j_ = Total number of individuals sampled

### Ethics statement

The study complies with ethical guidelines at Swansea University.

## Results

### Variation in body size and condition factor

The body size (total length) of topmouth gudgeon varied from 33 to 100 mm, and the fresh weight from 44 to 50 mg. Mixture analysis revealed that the vast majority of individuals (98%) were 1 year old (mean length = 56mm, SD = 12.3), and only three individuals (2%) were two years old (mean length = 95 mm, SD = 5.06). There were no significant differences among ponds in the length (*F*_3,110_ = 2.008, *P* = 0.117) or weight (*F*_3,110_ = 1.934, *P* = 0.1282) of one year old fish, but there was a significant difference in their condition factor (*F*_3,110_ = 5.1, *P* = 0.002; Figure 2) with fish in Turbine Pond having a better condition than those in Dyfatty pond (*P* =0.006) or the Upper Lliedi Reservoir (*P* =0.003).

**Figure 2.**
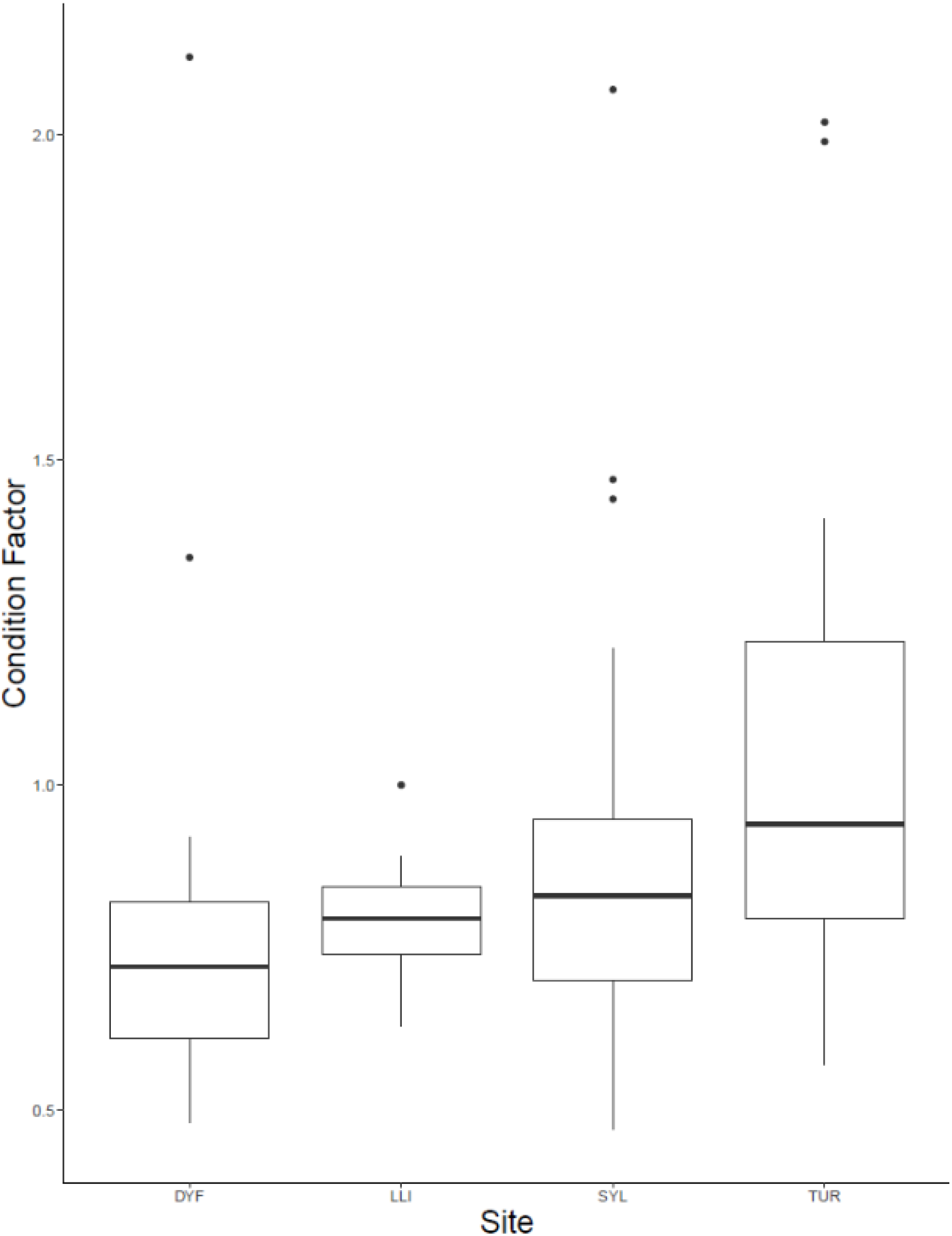
Variation in Fulton’s body condition factor of topmouth gudgeon across study sites.

### Diet analysis

The proportion of fish with empty stomachs differed significantly among sites (GLM binary logistic regression: deviance = 22.09, *P* <0.001), but was unrelated to body size (deviance = 2.57, *P* = 0.417), or condition factor (deviance = 0.14, *P* = 0.70). The pond with the greatest incidence of empty stomachs was DYF (21/25 or 84%), followed by SYL (19/29 or 65.5%), while TUR had the lowest incidence (6/27 or 22.2%). Tukey multiple comparisons at 95% family-wise confidence level show that the incidence of empty stomachs was significantly lower at TUR than at DYF (*P* <0.001) or SYL (*P* <0.005), while no significant difference was observed between DYF and SYL (*P* = 0.237).

The majority of stomach contents (756 food items or 67% of the total) was composed of zooplankton, including *Daphniidae* (268 specimens or 23.8%), *Bosminidae* (136 specimens or 12%), *Chydoridae* (348 specimens or 30.9%), and *Cyclopidae* (4 specimens or 0.3%). The remaining was composed of plant seeds (272 items or 24.1%), unidentified insect parts (12 items or 1%), and unidentified organic and plant material (88 items or 7.8%; Figure 3). Mean prey size varied considerably, being largest in insect parts (2.14mm), followed by *Cyclopidae* (1.51mm), *Daphniidae* (1.28mm), *Bosminidae* (0.67mm), plant seeds (0.62mm), and *Chydoridae* (0.48mm).

**Figure 3.** Individual variation in the diet of topmouth gudgeon (% items) across study sites showing Shannon diversity index (H).

Analysis of similarities (ANOSIM) indicated that the diet of TMG differed markedly among sites (R = 0.28, *P* = 0.005), and this was reflected in different prey diversity estimates. Thus, the diet of fish at DYF had a much lower prey diversity (mean bootstrapping estimate *H* = 0.148, 95CI = 0.089-0.218) than those at TUR (*H* = 1.270, 95CI = 1.223-1.317) or SYL (*H* = 1.244, 95CI = 1.173-1.318). Pairwise Hutcheson t-test comparisons indicated that there were statistical differences between TUR and DYF (*P* <0.001) and between SYL an DYF (*P* <0.001), but not between TUR and SYL (*P*=0.392).

Differences in diet were reflected in measures of diet breadth (Table 2). Thus, while fish at TUR and SYL had similarly wide diets (bootstrapping estimates B, TUR = 3.286, 95CI = 1.666-5.005; SYL = 3.270, 95CI = 1.611-4.998), the diet breadth at DYF was much narrower and more specialised (B = 1.522, CI =1.058- 2.000).

**Table 2.** Diet of topmouth gudgeon based on stomach content analysis (% number).

| Pond | *Daphniidae* | *Bosminidae* | *Chydoridae* | *Cyclopidae* | Insect parts | Unidentified items | Seeds | No. prey items | Percentage empty stomachs (%) | Levins's *B* (niche breadth) | Lower CI | Upper CI |
| --- | --- | --- | --- | --- | --- | --- | --- | --- | --- | --- | --- | --- |
| TUR | 0.38 | 0.16 | 0.39 | 0.01 | 0.02 | 0.04 | 0.00 | 664 | 22.2 | 3.07 | 1.666 | 5.005 |
| SYL | 0.09 | 0.15 | 0.43 | 0.00 | 0.00 | 0.33 | 0.00 | 184 | 65.5 | 3.07 | 1.611 | 4.998 |
| DYF | 0.00 | 0.00 | 0.03 | 0.00 | 0.00 | 0.00 | 0.97 | 280 | 84.0 | 1.06 | 1.058 | 2.000 |

### Stable Isotope Analysis

The carbon content in the muscle of topmouth gudgeon ranged between 41% and 60% among individuals, and differed significantly among ponds (Figure 4a, *F*_3,113_ = 11.54, *P* <0.001), being lowest in the LLI reservoir and highest in the eutrophic SYL pond (posthoc pairwise HSD, SYL-LLI, *P* <0.001; TUR-LLI, *P* <0.001). Carbon content did not depend on body length (t = −0.385, *P* =0.701), condition factor (t = −0.035, *P* =0.972) or the presence of food items in the stomach (t = −0.69, *P*=0.490). The nitrogen content varied from 12% to 18% among individuals, and as with %C, it also varied significantly among ponds (Figure 4b, *F*_3,113_ = 6.77, *P* <0.001), but did not depend on body length (t = 0.51, *P* = 0.610), condition factor (t = 0.31, *P* = 0.754), or presence of food items in the stomach (t = −1.51, *P* = 0.133). Nitrogen content was lowest in the reservoir and highest in the eutrophic angling pond (posthoc pairwise HSD, SYL-LLI *P*<0.001; SYL-DYF *P* = 0.004).

**Figure 4.**
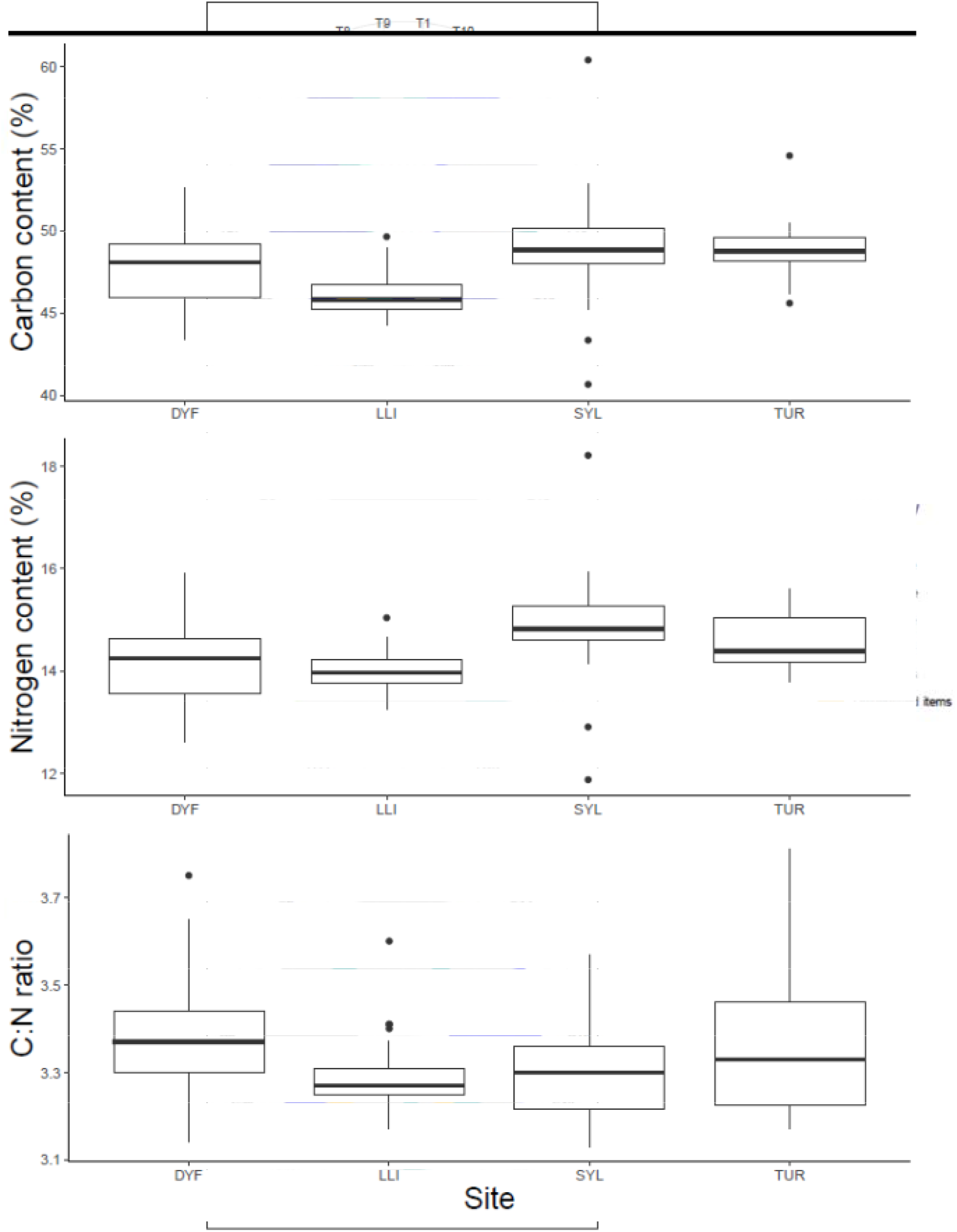
Variation in (a) % Carbon content, (b) % Nitrogen content and (c) C:N ratios of topmouth gudgeon across study sites.

C:N ratios (a measure of lipid reserves) varied between 3.13 and 3.81 among individuals, but were unaffected by body length (t = −1.22, *P* =0.224), condition factor (t = 0.2, *P* =0.835), or presence of food items in the stomach (t = −0.96, *P*=0.335). They varied significantly among sites (Figure 4c, *F*_3,113_ = 4.15, *P* = 0.007), being highest in one of the small ornamental ponds, and lowest in the reservoir (DYF-LLI, *P* = 0.016).

δ^13^C varied markedly among individuals (range = −30.95‰ to −16.43) and differed significantly among all sites (Figure 5; *F*_3,113_ = 24.77, *P* <0.001), being highest in the small ornamental ponds and lowest in the eutrophic angling pond (pairwise comparisons, LLI-DYF, *P* <0.001; TUR-DYF, *P* <0.001; SYL-LLI, *P* <0.001; TUR-SYL, *P* <0.001). δ^13^C was unaffected by body length (t = −0.09, *P* =0.927) or condition factor (t = −1.14, *P* =0.909), but increased (i.e. it was more enriched) among fish that had food in their stomachs (t = 2.90, *P*=0.004).

**Figure 5.**
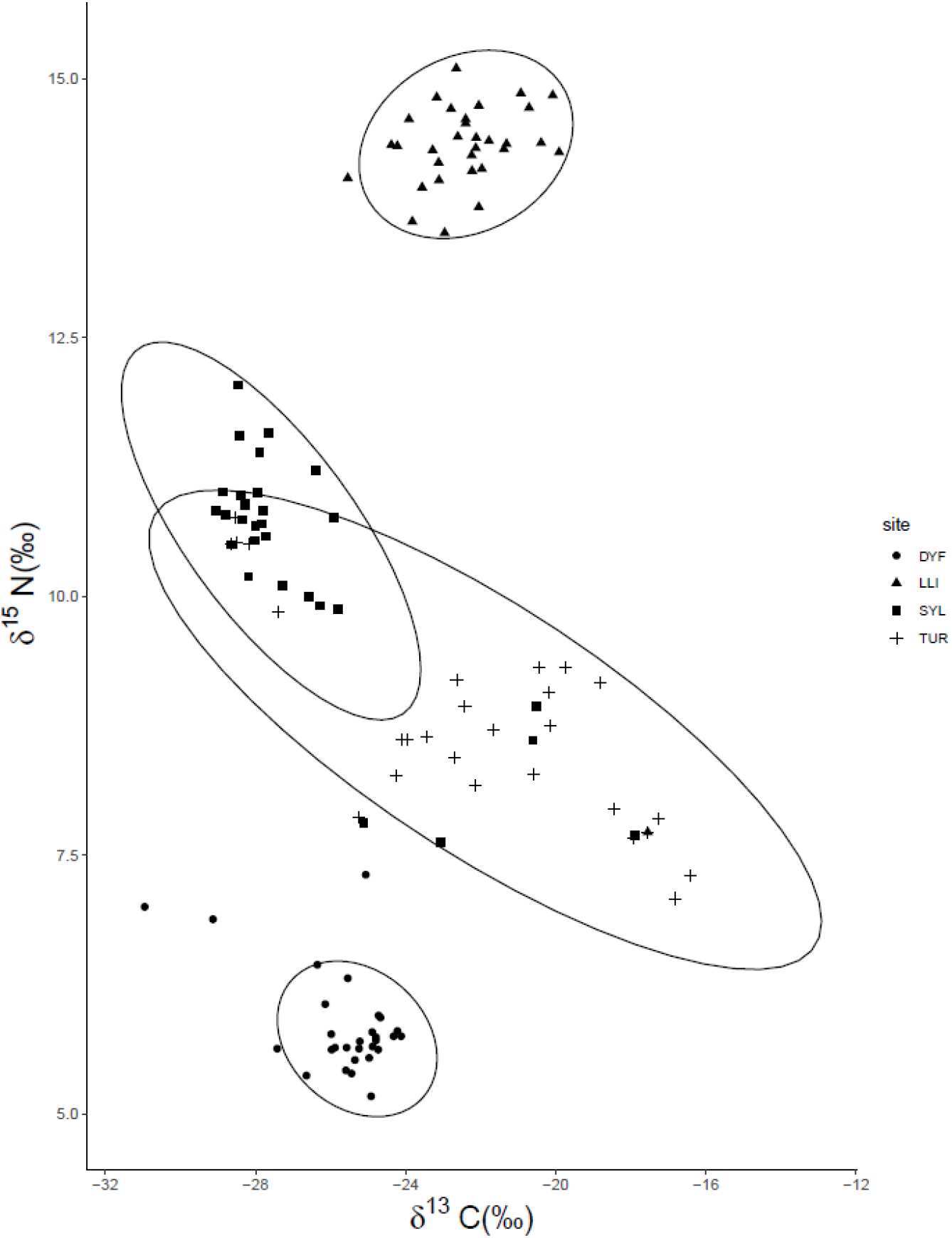
Biplot of δ13C vs δ15N of topmouth gudgeon (95% confidence ellipses) across study sites.

δ^15^N varied from 5.17‰ to 15.10‰ among individuals, and also varied significantly among all sites (Figure 5; *F*_3,113_ = 574.27, *P* <0.001), all ponds being significantly different from each other (*P* <0.001), being highest in the reservoir and lowest in the small ornamental ponds. As for δ^13^C, δ^15^N was unaffected by body length (t= −0.567, *P* =0.572) or condition factor (t = −0.156, *P* =0.876), but increased (i.e. it became more enriched) among fish that had food in their stomachs (t = −3.23, *P*=0.001).

Examination of the isotopic δ^13^C vs δ^15^N plot (Figure 5) indicate a substantial trophic discrimination among study sites, particularly with respect to δ^15^N, with the large, deep reservoir (LLI) being nitrogen enriched and the small ornamental pond (DYF) being nitrogen depleted compared to the other sites.

## Discussion

Our study sheds light on the topmouth gudgeon’s trophic plasticity and shows that its diet varies markedly among sites, even between neighbouring ponds only a few kilometres away from each other. Trophic plasticity, i.e. the ability to switch diets, is expected to facilitate establishment success among invasive species (Pettitt-Wade et al. 2015; Schröder, Garcia de Leaniz 2011), and may have allowed the topmouth gudgeon to colonise different environments, exploit novel food resources, and outcompete many native fish species (Záhorská et al. 2009; Záhorská et al. 2010). However, trophic plasticity also carries costs (Shea, Chesson 2002). Trophic shifts are common in response to changes in habitat and food availability (Werner, Hall 1977), but can also represent a strategy to avoid potential trophic competition (Jackson et al. 2012). In our study, one year old topmouth gudgeon with mean size of 56mm were feeding on relatively small food items, most of which were within the 0.5-0.6 mm size range, and hence approximately 1% of their body size. This is considerably lower than the median of 10-20% body size for the prey of many piscivorous freshwater fish (Gaeta et al. 2018). The consumption of small zooplankton, shown at all our sites, has been reported previously (Arnold 1990; Rosecchi et al. 1993), along with a diet composed of benthic crustaceans (Rosecchi et al. 1993; Xie et al. 2000), chironomids (Declerck et al. 2002), phytoplankton and algae (Muchacheva 1945). However, the reliance on plant seeds found at one of our sites was unexpected, and may have been the consequence of strong interspecific competition.

Results from the analysis of stomach contents were largely confirmed by results from SIA and seem to reflect the combined effects of food availability and competition. Thus, the widest variation in δ^13^C and δ^15^N was observed at one of the small recreational ponds (TUR), which is consistent with a generalist diet largely unaffected by inter-specific competition as only one other fish was recorded (European eel). In contrast, at the other recreational pond (DYF) topmouth gudgeon coexisted with four other species (eel, perch, rudd, and roach), displayed a very high occurrence of individuals with empty stomachs (84%) and relied almost exclusively on plant seeds. This may explain why they attained a much lower condition factor, and displayed a lower trophic level, suggesting that plant seeds are nutritionally poor and not a preferred food resource. Britton et al. (2010) have shown that roach occupies a similar trophic position than topmouth gudgeon and can result in competition for food, while rudd occupies a higher trophic level and may prey on topmouth gudgeon. Therefore, the presence of competitors and predators may have forced the topmouth gudgeon to adopt a more specialized feeding behaviour in this pond. In a similar way, the narrow trophic position of TMG at the reservoir could be explained by high predatory pressure, since this reservoir is managed by an angling association which regularly stocks it with trout for recreational fishing.

We found that the body condition factor (a measure of growth performance - Froese, 2006) and the C:N ratio (a proxy for stored lipids – Jardine et al. (2008)) was highest in the pond with the most diverse diet, widest diet breadth, and fewest competitors (Turbin pond where only he European eel was reported). This was also reflected in high carbon and nitrogen contents, as well as in high δ13C values. In aquatic food webs, high δ13C values tend to correlate with better feeding conditions and less competition (Hinz et al. 2017), which is consistent with our results. In contrast, TMG sampled in the much larger reservoir where predatory salmonids abound, had the lowest C:N ratios, indicative of low lipid reserves, as well as low carbon and nitrogen contents, which are also consistent with poor feeding conditions (Vander Zanden et al. 1999).

Comparisons of SIA values from our study and other studies (Table 3) indicate that TMG in Wales generally have higher δ^13^C and δ^15^N values (i.e more enriched) than those reported in another study in Great Britain (Britton et al. 2010), indicating a higher trophic level (δ^15^N) and a different primary source of carbon (δ^13^C). Our δ13C values are relatively high and closer to the ones reported in China - where topmouth gudgeon is a native species (Gao et al. 2017; Mao et al. 2012; Mao et al. 2016) than to values observed in Belgium where the species is invasive (Tran et al. 2015). High δ^13^C values are more typical of benthic food pathways, compared to pelagic food pathways which tend to be more ^13^C depleted (Pinnegar, Polunin 1999). Variation in δ^15^N was also greater in our study, and one of our sites (DYF) had the lowest δ^15^N value reported for the species; this coincides with a diet dominated by plant seeds. In general, the variability among sites is so high that for example δ^15^N values obtained from fish at one our sites (LLI) are more similar to those found in fish from Lake Taihu in China (Mao et al. 2012) than to those at neighbouring sites in Wales. This indicates that it is difficult to predict the diet or trophic position of TMG even within small geographical areas.

**Table 3.**
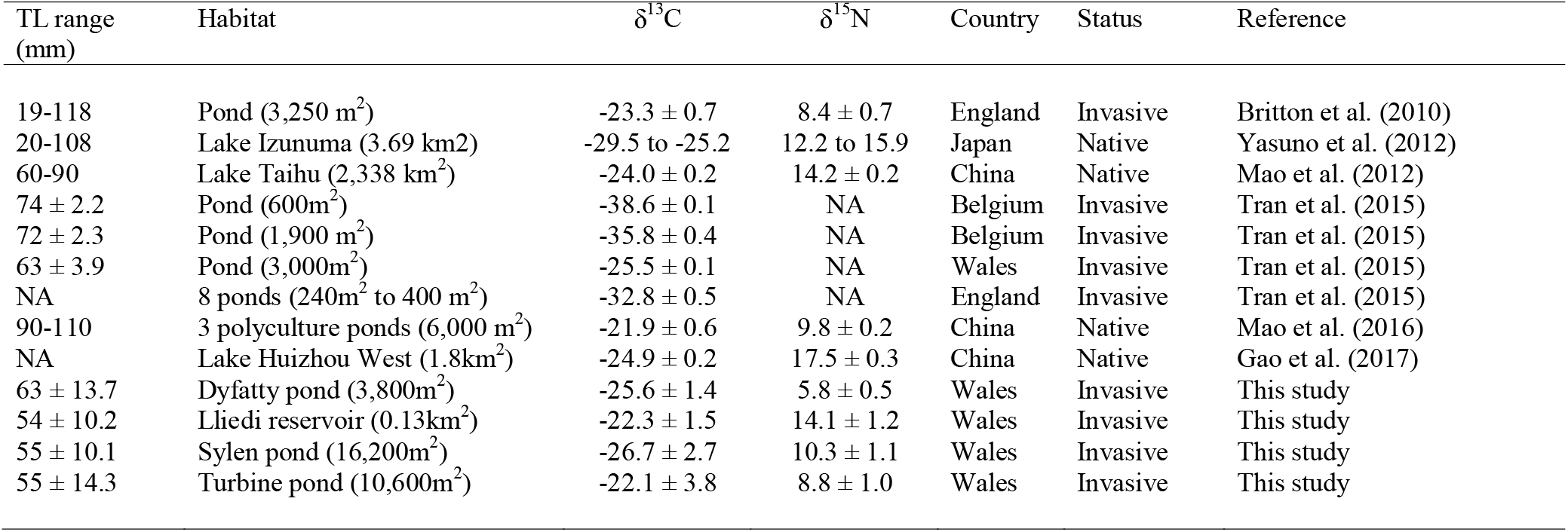
Comparison of isotope value for topmouth gudgeon across locations

| TL range<br>(mm) | Habitat | $\delta^{13}\text{C}$ | $\delta^{15}\text{N}$ | Country | Status | Reference |
| --- | --- | --- | --- | --- | --- | --- |
| 19-118 | Pond (3,250 m<sup>2</sup>) | -23.3 ± 0.7 | 8.4 ± 0.7 | England | Invasive | Britton et al. (2010) |
| 20-108 | Lake Izunuma (3.69 km<sup>2</sup>) | -29.5 to -25.2 | 12.2 to 15.9 | Japan | Native | Yasuno et al. (2012) |
| 60-90 | Lake Taihu (2,338 km<sup>2</sup>) | -24.0 ± 0.2 | 14.2 ± 0.2 | China | Native | Mao et al. (2012) |
| 74 ± 2.2 | Pond (600m<sup>2</sup>) | -38.6 ± 0.1 | NA | Belgium | Invasive | Tran et al. (2015) |
| 72 ± 2.3 | Pond (1,900 m<sup>2</sup>) | -35.8 ± 0.4 | NA | Belgium | Invasive | Tran et al. (2015) |
| 63 ± 3.9 | Pond (3,000m<sup>2</sup>) | -25.5 ± 0.1 | NA | Wales | Invasive | Tran et al. (2015) |
| NA | 8 ponds (240m<sup>2</sup> to 400 m<sup>2</sup>) | -32.8 ± 0.5 | NA | England | Invasive | Tran et al. (2015) |
| 90-110 | 3 polyculture ponds (6,000 m<sup>2</sup>) | -21.9 ± 0.6 | 9.8 ± 0.2 | China | Native | Mao et al. (2016) |
| NA | Lake Huizhou West (1.8km<sup>2</sup>) | -24.9 ± 0.2 | 17.5 ± 0.3 | China | Native | Gao et al. (2017) |
| 63 ± 13.7 | Dyfatty pond (3,800m<sup>2</sup>) | -25.6 ± 1.4 | 5.8 ± 0.5 | Wales | Invasive | This study |
| 54 ± 10.2 | Lliedi reservoir (0.13km<sup>2</sup>) | -22.3 ± 1.5 | 14.1 ± 1.2 | Wales | Invasive | This study |
| 55 ± 10.1 | Sylen pond (16,200m<sup>2</sup>) | -26.7 ± 2.7 | 10.3 ± 1.1 | Wales | Invasive | This study |
| 55 ± 14.3 | Turbine pond (10,600m<sup>2</sup>) | -22.1 ± 3.8 | 8.8 ± 1.0 | Wales | Invasive | This study |

## Conclusions

Our study indicates that when the topmouth gudgeon colonises a new area it may be difficult to predict its trophic impacts from local information, since differences in diet are largely independent of geographical proximity. Populations living in neighbouring water bodies can have diets more dissimilar than those from populations thousands of kilometres away. Given the importance diet information has on models that attempt to predict the risks posed by invasive fish, such as FISK (Copp 2013; Copp et al. 2009; Mastitsky et al. 2010; Simonovic et al. 2013), uncertainty on the expected diet of the topmouth gudgeon could result in incorrect risk assessments. To improve the precision of risk assessments, information on trophic level seems to be important for predicting the diet, and therefore the likely feeding impacts, of topmouth gudgeon on native communities. As trophic impacts will likely vary widely from site to site, our study shows that the use of SIA could be used, perhaps in combination with novel uses of eDNA for species detection (Robinson et al. 2019), to prioritize control and eradication measures that consider variation in diet.

## Acknowledgments

We are grateful to Ida Tavner and Emma Keenan (Natural Resources Wales) for providing information on the study sites and TMG samples, to Peter Jones (Swansea University) for help trapping topmouth gudgeon, and to Catherine Bradford and Therry Northam (Dwr Cymru Welsh Water) for permission to sample the Lliedi reservoir. Funding was provided by the EC Horizon 2020 Aquainvad-ED project (Marie Sklodowska-Curie ITN-2014-ETN-642197) to SC.

## Author Contributions & Competing Interests

CGL and SC designed the study and secured the samples and funding. MR collected the data and carried out the analyses with advice from CGL. MR and CGL wrote the paper with contributions from SC. The authors declare no competing interests.

